# Plasma, not extracellular vesicles, disrupts the blood-brain barrier in eclampsia

**DOI:** 10.1101/2025.03.03.641338

**Authors:** Jesenia Acurio, Felipe Troncoso, Estefanny Escudero-Guevara, Hermes Sandoval, Manu Vatish, Pablo Torres-Vergara, Lina Bergman, Carlos Escudero

**Author notes:** These two authors contribute equaly to this manuscript (Jesenia Acurio, Felipe Troncoso). Correspondence: Carlos Escudero, MD PhD Vascular Physiology Laboratory Group of Research and Innovation in Vascular Health Basic Sciences Department Faculty of Sciences Universidad del Bio-Bio Chillán, Chile /. **Financial disclosure**: Fondecyt 1240295. LB is found by Research grants from the Swedish Research Council, STINT, Märta Lundqvist stiftelse, Swedish Society of Medicine, SSMF, Jane and Dan Olssons stiftelse and Wallenberg Center for Molecular and Translational Medicine.

## Abstract

**Background:** Eclampsia is a severe complication of preeclampsia involving blood-brain barrier (BBB) disruption. While small extracellular vesicles (sEVs) contribute to endothelial dysfunction in preeclampsia, their role in eclampsia remains unclear. Magnesium sulfate (MgSO₄), the standard treatment, may mitigate BBB injury. We examined the effects of plasma and plasma-derived sEVs from women with eclampsia on BBB integrity and the potential modulatory role of MgSO₄.

**Methods:** Plasma and plasma-sEVs were isolated from women with normotensive pregnancies (n=18), preeclampsia (n=19), preeclampsia with organ complications (n=17), and eclampsia (n=20). An *in vitro* BBB model based on the culture of human brain endothelial cells was used to evaluate electrical resistance (TEER), Dextran 70 kDa permeability, and cytoskeletal alterations in the presence of women’s plasmas or plasma-sEVs. The uptake of fluorescently labeled sEVs in the absence or pretreatment (-3 h) with MgSO₄, and sEVs cargo of relevant proteins involved in BBB regulation, were analyzed.

**Results:** Plasma from women with eclampsia disrupted the BBB, with marked reductions in TEER and increased permeability compared to normotensive controls, preeclampsia, and preeclampsia with organ complications. In contrast, plasma-sEVs of women with eclampsia caused less BBB impairment than plasma-sEVs from normotensive controls or preeclampsia, correlating with reduced sEVs uptake by brain endothelial cells. Lower levels of eNOS and TNF-α in eclampsia-derived sEVs compared to normotensive controls were founnd. MgSO₄ treatment further diminished sEVs uptake.

**Conclusions:** Plasma, rather than sEVs, appears to drive BBB disruption in eclampsia. MgSO₄ may influence these effects by reducing sEVs uptake and altering their protein cargo.

## Introduction

Eclampsia, characterized by the onset of generalized seizures in women who experience preeclampsia, is associated with high maternal morbidity and mortality^1^. The condition is particularly severe in low-and middle income countries^2^. Despite MgSO₄ being the gold-standard prophylaxis and treatment for eclampsia^3,4^, the precise pathophysiological mechanisms underlying eclampsia and the mechanism of action of MgSO₄ remain poorly understood.

Recent advances remark on the potential role of circulating small extracellular vesicles (sEVs) as mediators of endothelial dysfunction during preeclampsia^5,6^. sEVs are lipid bilayer-enclosed particles released by cells, carrying bioactive molecules such as proteins, lipids, and nucleic acids reflective of their cell of origin. Within pregnancy, the placenta actively releases sEVs into the maternal circulation^7,8^, serving as vehicles for maternal-fetal communication. However, in pathological conditions such as preeclampsia, sEVs become potent drivers of systemic vascular inflammation and endothelial injury^9–11^. Our group has shown that sEVs derived from the plasma of women with preeclampsia decrease the protein level of claudin 5 (CLDN5), leading to increased blood brain barrier (BBB) permeability^12^. The effect of sEVs in eclampsia has not been explored.

Magnesium sulfate (MgSO4) has demonstrated stabilizing the BBB in experimental models potentially through its anti-inflammatory properties, calcium blockade, and beneficial effects on endothelial cell integrity^13,14^. Moreover, MgSO4 prevented BBB disruption induced by circulating or placenta-derived sEVs in our *in vitro* and *in vivo* systems^14^. Whether MgSO₄ modulates the interaction between circulating sEVs and the cerebrovascular endothelium during eclampsia remains unknown. Such insights could have profound implications for optimizing therapeutic strategies and mitigating the severe neurological outcomes of this condition.

Therefore, this study aimed to elucidate whether plasma and plasma-derived sEVs from women with eclampsia can generate BBB dysfunction. We hypothesized that plasma and/or sEVs from women with eclampsia would lead to increased BBB permeability through alterations in brain endothelial cell integrity. Additionally, we investigated whether MgSO₄ treatment modulates sEVs uptake by brain endothelial cells and influences BBB integrity.

## Methods

### Population

Women participating in the PROVE biobank at Tygerberg Hospital, South Africa, were eligible for inclusion in the study^15^. Participants were included between 12/04/2018 until 31/01/2020. All variables were prospectively entered by research midwives, obtained from interviews or extracted from the medical charts, and double-checked for accuracy. All women were managed according to clinical routine including administration of MgSO_4_ in case of threatening preterm birth <32 weeks and for seizure prophylaxis in preeclampsia.

### Exposures

Exposures were preeclampsia, preeclampsia with organ complications without cerebral features, and eclampsia. Diagnosis of preeclampsia and eclampsia were defined using the ISSHP classification^16^. In addition, significant proteinuria was required for diagnosis of preeclampsia (2+ protein on a dipstick and/or urine protein/creatinine ratio above 30 mg/mmol). Preeclampsia with organ complications was defined according to a core outcome set of preeclampsia-related complications^17^.

### Control

Normotensive controls were defined as women with a pregnancy where blood pressure did not exceed 140/90 mm Hg. Exclusion criteria for exposure and control groups were pre-existing neurological disorders and cardiovascular disease.

The research was performed by the principles expressed in the Declaration of Helsinki and under the authorization of the respective Ethical Review Boards. Ethical approval for the inclusion of women for this study was obtained from the Stellenbosch University Health Research Ethics Committee (protocol numer N18/03/034; Federal Wide Assirance number 00001372; institutional review board number: IRB0005239). All participants gave their informed consent before sample collection.

### Outcomes

#### TEER and Permeability

To analyze the function of the BBB *in vitro*, we used a previously validated Transwell^®^ system using the hCMEC/D3 cell line^18–20^. Briefly, cell monolayers were exposed to plasma (1:10 v/v, 12 h) or sEVs (100 μg sEVs per Transwell, 12 h) belonging to the respective experimental groups (see below). Measurements of transendothelial electrical resistance (TEER) (EVOM2, World Precision Instruments, USA) and cell monolayer permeability to high molecular weight fluorescent dye (Dextran 70 kDa fluorescein-5-isothiocyanate-FITC) were performed as described previously^18,19^.

### F-actin disorganization

F-Actin fibers were identified using phalloidin iFluor 488 (1:500 v/v dilution) (Abcam, Cambridge, UK; ab176753,) in hCMEC/D3 cells cultured over a glass put in 24-well plates and treated as described above using plasma or plasma-sEVs for 6 hours. After overnight incubation at 4°C, samples were consecutively washed with 1 X phosphate buffer saline (PBS) solution. Then, nuclear labeling (DAPI) was added in dilution (1:10,000 v/v) for 20 minutes. Finally, all samples (i.e., cultured over the glass) were transferred onto a 26 x 76 mm slide (Knittel Glass, BS, DE) and covered by a glass coverslip supported by mounting fluid for subsequent visualization using fluorescent microscopy (Motic model BA410, Motic, HK, China). Images were captured with 100 X magnification and a resolution of 2580 x 1944 pixels per image. Three RGB chromatic channels (Red, Green, and Blue) were used, with their corresponding filters (FITC) (Ex 330-380), (TRITC) (Ex 450-490), and (DAPI) (Ex 510-560). Then, captured fluorescent cell images were analyzed with the Image J program (National Institutes of Health, NIH) using the Mexican Hat filter in each analysis, which preserves high frequencies, thus highlighting the fibers. The Ride Detection plugins were applied to analyze the number and length of fibers. The length of the fibers was normalized by the area (i.e., pixels) of the respective cell.

### Biological samples

EDTA plasma from pregnant controls (n=18), preeclampsia (n=19), preeclampsia with organ complications without cerebral complications (n=17), and eclampsia (n=20) were obtained from the Preeclampsia Obstetric Adverse Events Biobank at the Tygerberg Obstetric Critical Care Unit in South Africa^15^ and used for all *in vitro* experiments. For *in vitro* experiments, 1% plasma was diluted in culture medium without fetal calf serum (v/v), and cells were exposed to each condition under varying treatment times, depending on the analysis performed.

### Cell viability and apoptosis assays

To analyze the effect of plasmas on cell viability, we used the CellTiter 96 Non-Radioactive Kit (Lot: 0000105232, Promega, Madison, WI, USA), following manufacturer instructions. hCMEC/D3 cells (Merck Millipore, Darmstadt, Germany) were cultured on a 96-well plate and, after reaching 80% confluence, were treated with plasma (24 h, 1:10 v/v per well) from respective experimental groups. Absorbance was analyzed using an Epoch spectrophotometer equipment (BioTek Instruments, VT, USA), set up at 570 nm^14^. Furthermore, we also measured caspase 3/7 activity using NucView 488 kit (Biotium, Germany) in hCMEC/D3 cells exposed to each condition (12 h, plasma 1:10 v/v per well) following manufacturer instructions^21^. Fluorescence was read on a plate reader at settings to 488 nm (excitation) and 520 nm (emission). This assay was complemented by analysis of BAX (Santa Cruz Biotechnology, Dallas, TX, USA; sc-7480, TX, USA) and Bcl-2 (Santa Cruz Biotechnology; sc-7382) proteins by Western blot. BAX/BCL2 ratio was estimated using the ImageJ 1.48 software (National Institute of Health, USA).

### Plasma sEVs Isolation, characterization, and protein content

EDTA-plasma sEVs were isolated following differential centrifugation and microfiltration protocol, as described previously^12,22^. Sequential centrifugation of plasma diluted in 1X PBS (pH 7,4) was performed.^23^. Briefly, after collecting plasma, we performed sequential centrifugation steps: (1) 300 *x g* for 10 minutes, (2) 2000 *x g* for 10 minutes, (3) 10,000 *x g* for 30 minutes, and (4) 120,000 *x g* for 2 hours at 4°C. The final supernatant was passed through a 0.22 μm filter and then centrifuged at room temperature at 120,000 *x g* for 18 hours. We recovered the pellet containing sEVs, resuspended it in PBS (pH 7.4), and passed it through a 0.22 μm filter. Finally, we performed the last centrifugation at 120,000 *x g* for 3 hours at 4°C. The pellet was resuspended in PBS (pH 7.4, previously depleted of sEVs), and passed it through a 0.22 μm filter.

sEVs were characterized by size and concentration using the NanoSight NS300 Instrument (Malvern Instruments, Ltd, Malvern, United Kingdom)^24^. Samples were analyzed in a liquid suspension (PBS pH 7.4, 1:1000 dilution) at room temperature. Measurements ensured that there were between 20 and 100 particles per frame. Negative controls had less than seven particles per frame. Each sample was measured in triplicate with the same camera setup, and Data was analyzed using NTA version 3.2 Dev Build 3.2.16 analytical software.

In addition, transmission electron microscopy (TEM) was performed in representative randomly chosen samples from women with normotensive pregnancies and preeclampsia, as indicated previously^12^. Ten microliters of each sEVs sample (in 1X PBS) were placed on formvar-carbon-coated copper grids. Samples were fixed with 4% paraformaldehyde (1:1, 20 min), treated (6 min) with 1% glutaraldehyde, and washed with molecular-grade water. For contrast, grids were treated (5 min) with 0.5% uranyl oxalate (pH 7.0), dried at room temperature, and imaged at 40,000 X to 80,000 X magnification using a Jeol Jem 1200 EXII TEM with a Gatan 782 camera (5Å resolution, 80 kV).

sEVs were also characterized in terms of protein markers using Western blot and the following Santa Cruz Biotechnology antibodies (Dallas, TX, USA): Alix, sc-53540; CD63, sc-5275; CD81, sc-7637; TSG101, sc-7964; and HSP70; sc-66048. Also, placental alkaline phosphatase (PLAP, sc-53414) as the placental marker was analyzed^12,14^.

### Protein content of BBB modulators in the plasma sEVs

Pools of plasma sEVs protein extractions were used to analyze the presence of potential BBB modulators, including the proinflammatory cytokine tumor necrosis factor-alpha (TNF-α, Santa Cruz Biotechnology, sc-52746); angiogenic modulators, including vascular endothelial growth factor (VEGF, Santa Cruz Biotechnology, sc-7269), placental growth factor (PlGF, Santa Cruz Biotechnology, sc-53414), and endothelial nitric oxide synthase (eNOS, BD Transduction Laboratories, Becton, NJ, USA; 610299), by Western blot.

For Western blot, total protein quantification was performed using Pierce ^TM^ BCA protein assay kit following manufacturer instructions (Thermo Fisher Scientific, Waltham, MA, USA). 50 μg of sEVs protein were separated in a 10% SDS-PAGE gel, transferred to a nitrocellulose membrane, and incubated with the respective primary antibodies. Ponceau red staining was used as a protein loading control.

### sEVs uptake by hCMEC/d3 cells and the effect of magnesium sulfate

To characterize the uptake of sEVs by hCMEC/d3 cells, the former were labelled with PKH67 Green Fluorescent Cell Linker Mini Kit (MINI67-1KT, Sigma-Aldrich, St. Louis, MO, USA), according the manufacturer protocol and cleared of free dye through an Amicon® Ultra Centrifugal Filter unit -100 kDa MWCO (Merck; Darmstadt, Germany). Briefly, 100 µg/ml of labelled sEVs were applied to hCMEC/D3 monolayers for 1 hour at 37°C. Cells were then fixed with paraformaldehyde (PFA 4% in PBS 1x) for 20 minutes at room temperature. Over several washing steps with PBS 1x, slides were incubated with DAPI stain for 20 minutes and mounted over microscopic slides using DAKO mounting medium. After this, slides were kept at 4°C until analysis by fluorescent microscopy (40X magnification) (Motic Scientific, San Antonio, TX, USA). To quantify the uptake of sEVs by hCMEC/D3 cells, the presence of green dots (i.e., sEVs) at the FITC channel was measured.

In parallel experiments, to visualize the effect of sEVs on cell uptake, we pretreated (-3 hours) the cells with magnesium sulfate (80 mg/L w/v) before adding sEVs, as we previously reported^14^.

### Statistics

All statistical analysis was performed using Prism GraphPad (V10, GraphPad Software, La Jolla, CA, USA). The Shapiro-Wilk normality test was used to discriminate between parametric and non-parametric distributions. The unpaired t-test (parametric), or Kruskal-Walli’s test (non-parametric), followed by an uncorrected Dunn’s test, was used accordingly. Values are expressed in the median with the interquartile range. A p-value of 0.05 was set as statistically significant. All experiments were performed in duplicate.

## Results

### Background characteristics

The baseline characteristics of the study participants are summarized in Table 1. Statistical testing was not performed for the baseline characteristics table, as the study was not designed or powered to detect differences in these variables. However, women with eclampsia were younger than those with preeclampsia with organ complications. Gestational age at delivery was lower in women with eclampsia compared to normotensive controls. Women in the preeclampsia with organ complications group exhibited a higher body mass index (BMI) compared to normotensive pregnancies. Severe hypertension and pulmonary edema were more prevalent in the preeclampsia with complications group (77% in both cases). Adverse perinatal outcomes, including intrauterine and neonatal death, were observed in 10 cases, predominantly among women with eclampsia and preeclampsia with complications.

**Table 1.** Characteristics of included women Figure legends.

|  | Eclampsia | Preeclampsia with complications | Preeclampsia | Normotensive Controls |
| --- | --- | --- | --- | --- |
|  | n=20 | n=17 | n=19 | n=18 |
| **Baseline characteristics** |  |  |  |  |
| Maternal age, years ± SD | 23.7 ± 5.2 | 30.6 ± 5.8 | 26.9 ± 5.5 | 27.5 ± 7.0 |
| Nulliparous, n (%) | 13 (65) | 5 (29) | 10 (53) | 6 (33) |
| Chronic hypertension, n (%) | 3 (15) | 4 (25) | 2 (10.5) | 0 (0) |
| BMI (kg/m<sup>2</sup>), mean ± SD | 25.8 ± 4.6 | 32.1 ± 9.5 | 28.5 ± 7.4 | 25.2 ± 6.8 |
| **At inclusion** |  |  |  |  |
| Magnesium sulfate, n (%) | 20 (100) | 17 (100) | 18 (95) | 2 (11) |
| **At birth** |  |  |  |  |
| Gestational age at birth (week + days) | 31+2 (4+6) | 31+6 (3+6) | 32+1 (4+2) | 35+2 (4+2) |
| **Mode of birth** |  |  |  |  |
| Vaginal n (%) | 7 (35) | 2 (12) | 6 (32) | 5 (28) |
| Planned caesarian, n (%) | 0 (0) | 1 (6) | 3 (16) | 11 (61) |
| Emergency caesarian, n (%) | 13 (65) | 14 (82) | 10 (52) | 2 (11) |
| Birthweight, gr, mean ± SD | 1594 ± 845.0 | 1636.7 ± 881.5 | 1666.8 ± 975.0 | 2522.5 ± 997.9 |
| Sex of newborn female, n (%) | 11 (55) | 9 (53) | 7 (37) | 5 (28) |
| Placental weight, g, mean ± SD | 363.8 ± 177.2 | 393.7 ± 159.3 | 381.6 ± 174.4 | 555.7 ± 158.6 |
| **Complications** |  |  |  |  |
| Recurrent eclampsia, n (%) | 6 (30) | N/A | N/A | N/A |
| Severe hypertension, n (%) | 11 (55) | 13 (77) | 6 (32) | 0 (0) |
| Stroke, n (%) | 1 (5) | 0 (0) | 0 (0) | 0 (0) |
| Cortical blindness, n (%) | 1 (5) | 2 (12) | 0 (0) | 0 (0) |
| Creatinine > 120, n (%) | 3 (15) | 4 (24) | 0 (0) | 0 (0) |
| Dialysis, n (%) | 0 (0) | 1 (6) | 0 (0) | 0 (0) |
| Pulmonary edema, n (%) | 1 (5) | 13 (77) | 0 (0) | 0 (0) |
| Inotropic support, n (%) | 0 (0) | 0 (0) | 0 (0) | 0 (0) |
| HELLP, n (%) | 6 (30) | 4 (24) | 0 (0) | 0 (0) |
| Postpartum hemorrhage, n (%) | 1 (10) | 2 (12) | 0 (0) | 1 (6) |
| GCS <13 | 7 (35) | 0 (0) | 0 (0) | 0 (0) |
| Intrauterine fetal death, n (%) | 3 (15) | 2 (12) | 2 (11) | 0 (0) |
| Neonatal death, n (%) | 1 (5) | 2 (12) | 0 (0) | 0 (0) |
| Placental abruptio, n (%) | 1 (5) | 2 (12) | 0 (0) | 1 (6) |

Glasgow coma scale (GCS). SD standard deviation

### Effect of plasma on the BBB *in vitro* model

Initially, we assessed the effects of plasma from women with normotensive pregnancies, preeclampsia, preeclampsia with organ complications, and eclampsia on cell viability (Figure 1A) or induced apoptosis (Figure 1B, 1C). Results showed that all plasmas significantly increased cell viability compared to basal conditions (i.e., cells without stimulation, Kruskal-Wallis test, p<0.0001). However, the magnitude of the cell viability enhancement was higher in preeclampsia compared with preeclampsia with organ complications or eclampsia (p=0.029 and p=0.010, respectively). Conversely, no significant changes in the caspase-3 activity (Figure 1B) of the Bax/Bcl2 ratio (Figure 1C) among the eclampsia/preeclampsia groups compared to normal pregnancy were found. Although, caspase-3 activity was higher in preeclampsia than preeclampsia with organ complications.

**Figure 1.**
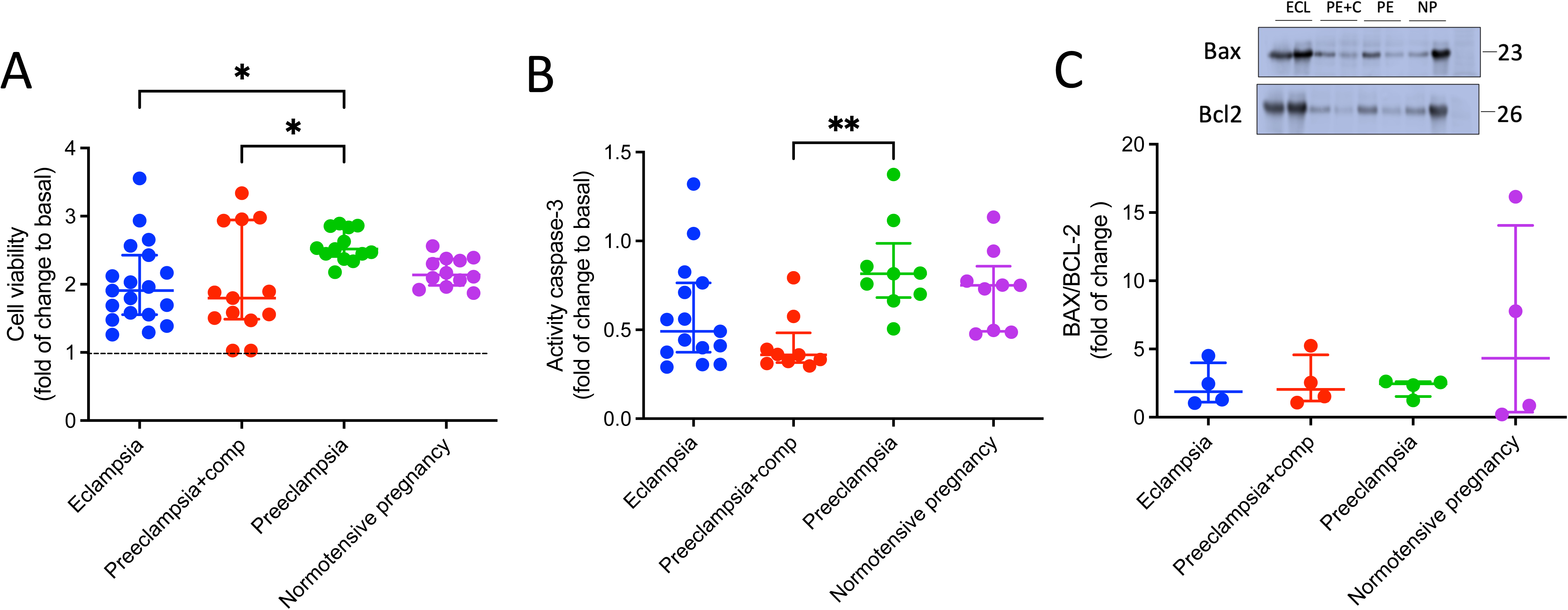
Effects of plasma of women with preeclampsia or eclampsia on hCMEC/D3 cells viability and apoptosis. **A)** Cell viability measured by MTT assay, **B)** Caspase 3 activity, and **C)** BAX/BCL-2 ratio (see Methods) in brain endothelial cells (hCMEC/D3) exposed (24 h) to plasma of women with eclampsia (ECL), preeclampsia with (PE+C) or without (PE) organ complications and normotensive controls (NP). Every dot represents an individual subject. Vales are expressed in the median with interquartile range. In A, line represent basal conditions without plasma estimulation. *p<0.05. **p=0.0040. Kruskal-Walli’s test, followed by Dunn’s multiple comparisons test.

Then, we analyzed whether plasmas of women with preeclampsia/eclampsia lead to impairment of the BBB *in vitro* model. Our results indicate that plasma from women with eclampsia disrupted the BBB model compared with the other three groups, as evidenced by a higher drop in the TEER values (Figure 2A, Kruskal-Wallis test, p=0.0003), and higher 70 kDa Dextran permeability assay (Figure 2B, Kruskal-Wallis test, p=0.049). In addition, increased alterations in the cell cytoskeleton demonstrated by reduced length of F-actin per cell was observed in the preeclampsia with complications group compared with normal pregnancy, preeclampsia and eclampsia groups (Figure 2C Kruskal-Wallis test, p=0.076).

**Figure 2.**
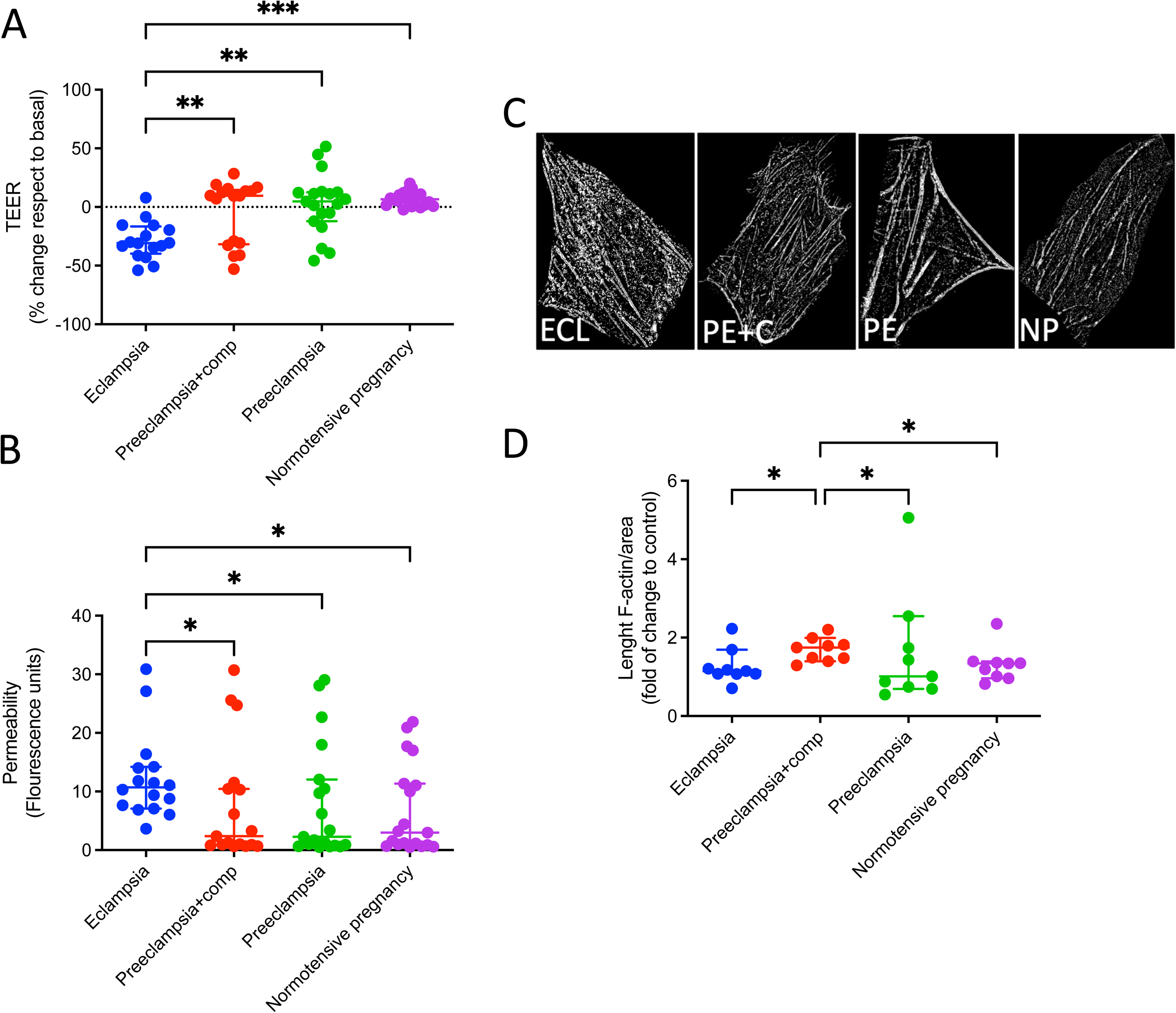
Effect of plasma of women with preeclampsia or eclampsia on the BBB *in vitro* model. **A)** Trans-endothelial electrical resistance (TEER) and **B)** Dextran 70 kDa permeability were analyzed in hCMEC/D3 cultures exposed (12 h) to plasma of women with eclampsia (ECL), preeclampsia with (PE+C) or without (PE) organ complications and normotensive controls (NP). **C)** Representative images of F-actin filaments in cells treated as in A. **D)** Estimation of F-actin length per cells, as an indicative of cytoskeleton disorganization. Every dot represents an individual subject. Vales are expressed in the median with interquartile range. *p<0.05. **p=0.0038; ***P=0.0005. Kruskal-Walli’s test, followed by Dunn’s multiple comparisons test.

### Effect of the plasma-derived sEVs on the *in vitro* BBB model

sEVs were successfully isolated from all study groups and characterized by nanoparticle tracking analysis, electron microscopy, and Western blotting for exosomal markers (Figure 3). No significant differences were observed in sEVs size (Figure 3A, 3C) or concentration (Figure 3A, 3D) across groups. Neither in the relative level of placental marker, PLAP (Figure 3E).

**Figure 3.**
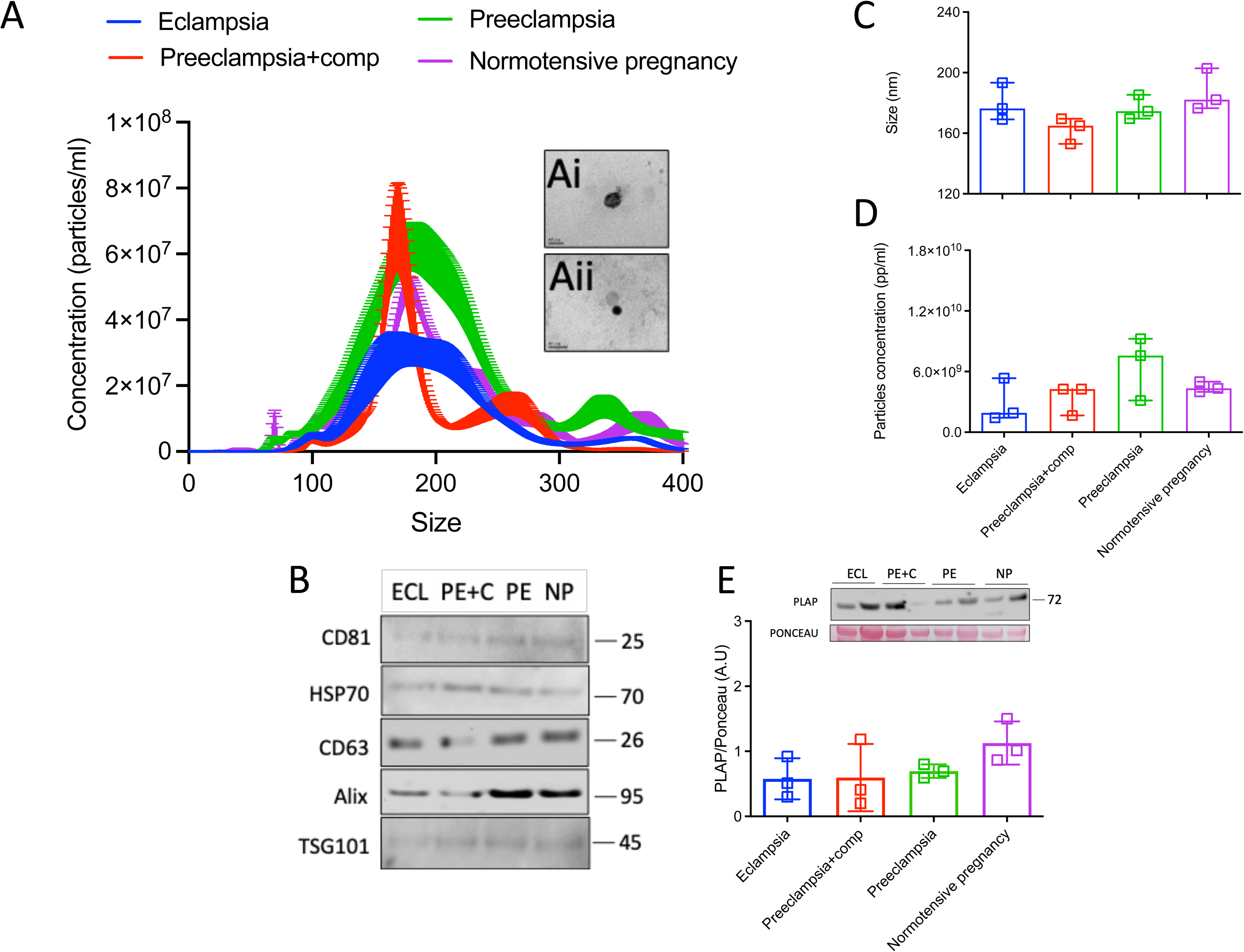
Characterization of sEVs isolated from plasma of women with preeclampsia or eclampsia. sEVs were isolated from the plasma of women with eclampsia (ECL), preeclampsia with (PE+C) or without (PE) organ complications and normotensive controls (NP) using the ultracentrifugation and filtration method (see Methods). **A)** Chart showing size and concentration distribution of sEVs from the experimental groups. Inserted images are representative sEVs observed in transmission electron microscopy (TEM) from ramdomly chosen normotensive pregnancy (Ai) or preeclampsia (Aii) groups. **B)** Representative images of sEVs marker (HSP70, CD81, Alix, TSG101, CD63). **C)** Size and **D)** Total number of sEVs from experimental groups. **E)** Representative image of placental marker (PLAP). The loading control was Ponceau Red staining. And densitometric analyses of PLAP/Ponceau ratio as indicated in E. Every dot represents an individual subject. Vales are expressed in the median with interquartile range.

Unlike plasma (Figure 2A-B), plasma-derived sEVs from women with eclampsia caused a milder reduction in TEER (p=0.0095) and a smaller increase in permeability (p=0.0075) compared to sEVs from women with normotensive pregnancy and women with preeclampsia (Figure 4A-B). F-actin fiber length was less affected by sEVs from women with eclampsia compared to preeclampsia-derived sEVs (Figure 4C-D, p=0.0049). These findings suggest that plasma, rather than sEVs, is the primary driver of BBB disruption in eclampsia.

**Figure 4.**
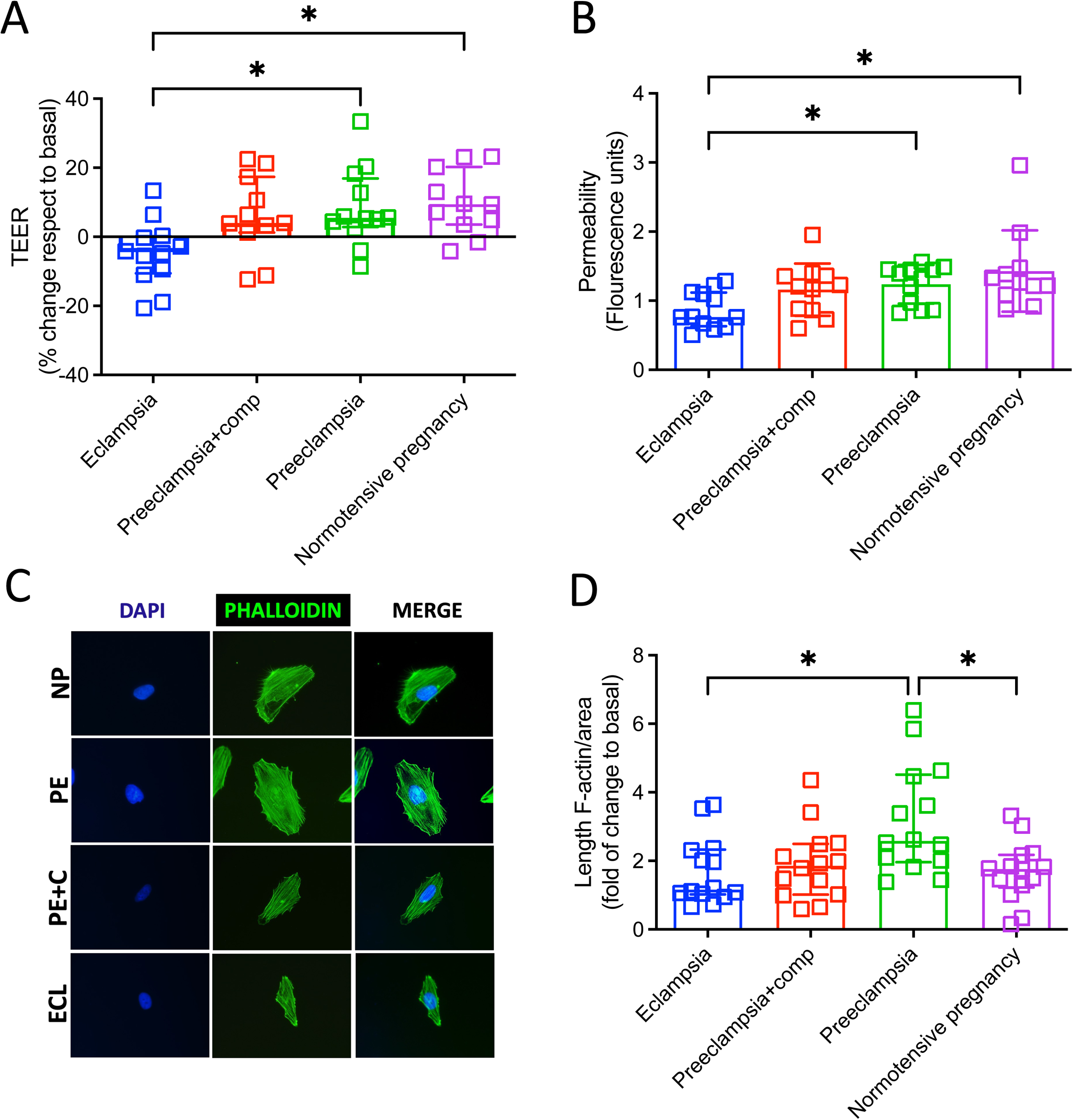
Effect of sEVs isolated from women with preeclampsia or eclampsia on the BBB *in vitro* model. sEVs isolated from the plasma of women with eclampsia (ECL), preeclampsia with (PE+C) or without (PE) organ complications and normotensive controls (NP) were used to analyze **A)** Trans-endothelial electrical resistance (TEER) and **B)** Dextran 70 kDa permeability in hCMEC/D3 cell monolayers after treatment (12 h). **C)** Representative images of F-actin filaments in cells treated as in A. **D)** Estimation of F-actin length per area as indicative of cytoskeleton disorganization. Every dot represents an individual subject, except in D, where every dot represents an individual cell (n=3 independent sEVs extractions were used per group). Vales are expressed in the median with interquartile range. *p<0.05. Kruskal-Walli’s test, followed by Dunn’s multiple comparisons test.

### Effect of magnesium sulfate on BBB uptake of sEVs

To investigate whether MgSO₄ treatment influenced sEVs uptake, we analyzed the percentage of hCMEC/D3 cells incorporating fluorescently labeled sEVs. Pretreatment with MgSO₄ (-3 h) reduced sEVs uptake in preeclampsia and normotensive groups (Figure 5A-B, p=0.0028). In the eclampsia group, sEVs uptake was already lower compared to normotensive and preeclampsia with complication groups (p=0.016 and 0.048, respectively, Figure 5C-D). Contrary to the other three groups, MgSO₄ treatment *in vitro* did not reduce uptake in the group of women with eclampsia, suggesting a possible preconditioning effect of MgSO₄ on circulating sEVs (Figure 5E).

**Figure 5.**
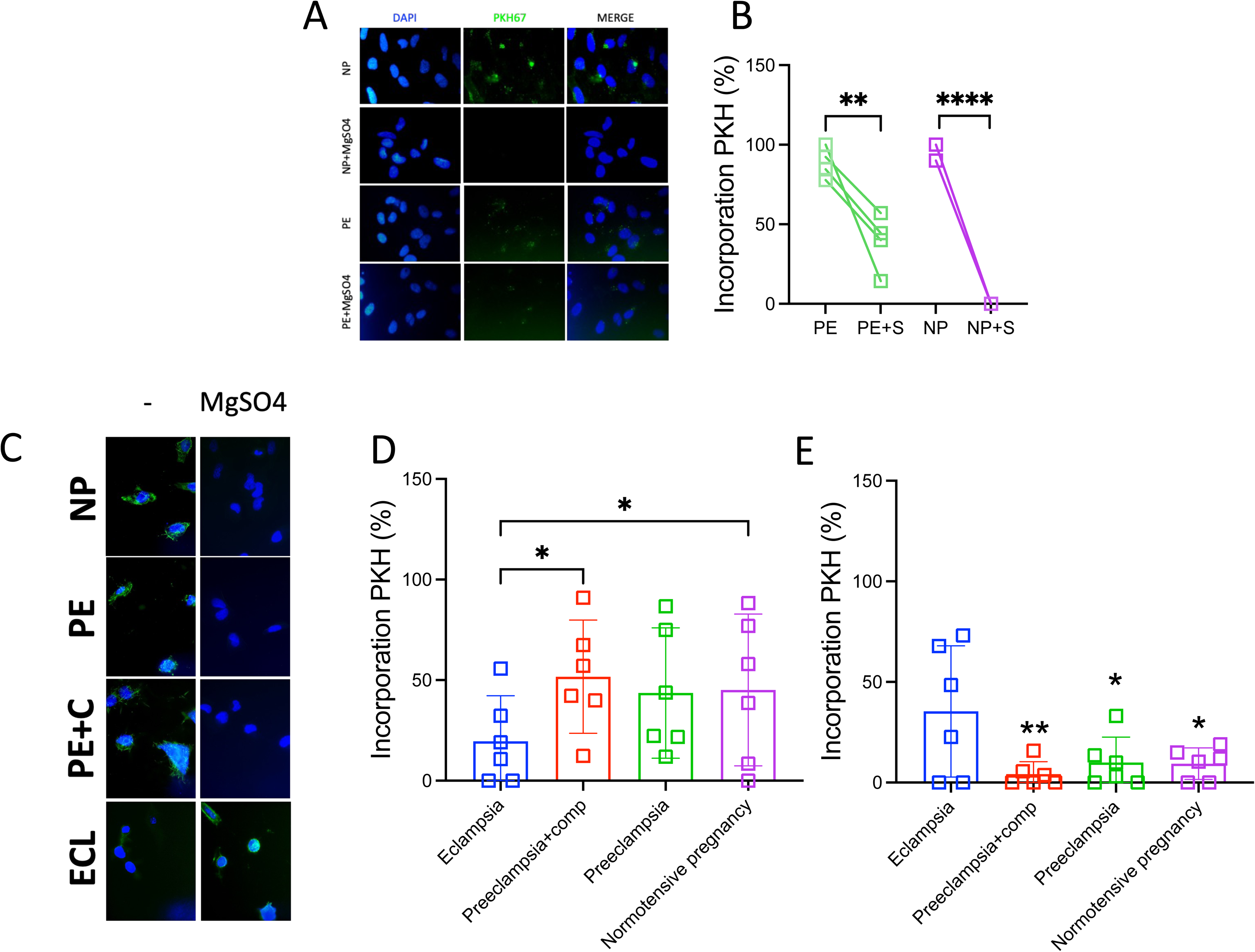
Effect of magnesium sulfate into sEVs uptake. **A)** Representative images of hCMEC/D3 cultures treated with sEVs isolated from plasma of women with normotensive pregnancies (NP) or preeclampsia in basal conditions (i.e., without any additional treatment) or pretreated (-3 h) with magnesium sulfate (MgSO4, +S). sEVs were treated with a fluorescent dye (PKH67) as indicated in Methods. **B)** Estimation of the percentage of cells that uptake sEVs (i.e., PKH67 positive) concerning the group of cells observed in each visual camp. **C)** Representative images of hCMEC/D3 cells pretreated or not with magnesium sulfate in all experimental groups (eclampsia (ECL), preeclampsia with (PE+C) or without (PE) organ complications and normotensive controls (NP)). **D)** Percentage of cells that uptake sEVs in cells treated with sEVs. **E)** Percentage of cells that uptake sEVs in cells pretreated (-3h) with magnesium sulfate and then treated with sEVs. Vales are expressed in the median with interquartile range. In B **p=0.0028; ***p=<0.001 Unpaired t-test. In D *p<0.05. Kruskal-Walli’s test, followed by Dunn’s multiple comparisons test. In E **p=0.018 or *p=0.033 Unpaired t-test versus respective group without MgSO4.

### Content of sEVs

Western blot analysis of plasma-derived sEVs revealed lower levels of endothelial nitric oxide synthase (eNOS) and tumor necrosis factor-alpha (TNF-α) in eclampsia-derived sEVs compared to normotensive controls (p=0.022 and p=0.0023, respectively, Figure 6A-G). No significant differences were observed in vascular endothelial growth factor (VEGF) or placental growth factor (PlGF) levels between groups. The reduced cargo of key endothelial regulatory proteins may contribute to the attenuated effect of eclampsia-derived sEVs on BBB integrity.

**Figure 6.**
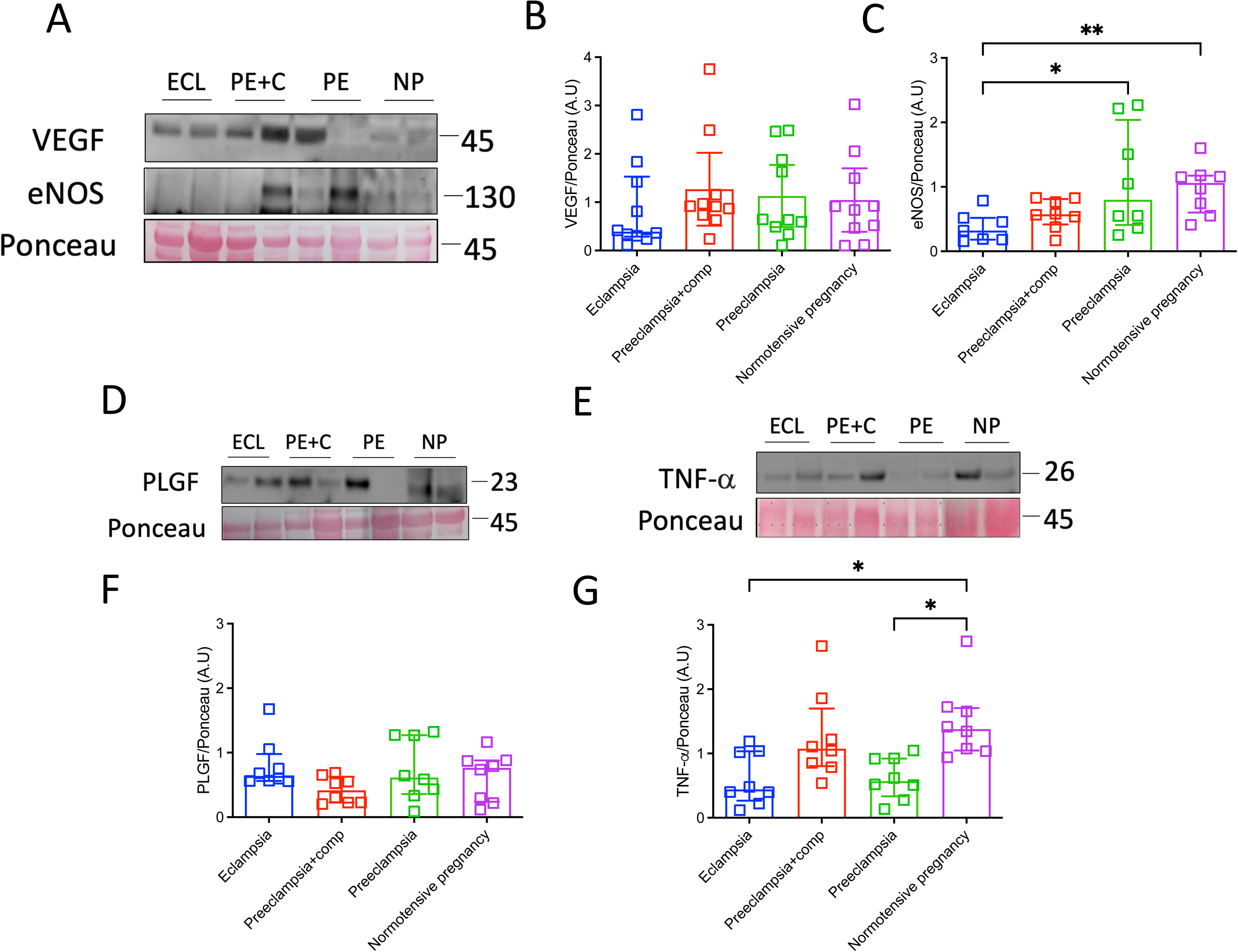
Analysis of cargo of sEVs of BBB modulatory proteins. **A)** Representative images of Western blot for VEGF and eNOS **D)** PlGF and **E)** TNF-α present in sEVs isolated from plasma of women with eclampsia (ECL), preeclampsia with (PE+C) or without (PE) organ complications and normotensive controls (NP). Ponceau staining was used for loading cargo. **B), C), F), G)** Densitometric analysis of respective analyzed proteins. Vales are expressed in the median with interquartile range. *P<0.05. **P=0.0037. Kruskal-Walli’s test followed by Uncorrected Dunńs test.

## Discussion

Our findings demonstrate that plasma from women with eclampsia significantly disrupts BBB integrity, as evidenced by reduced TEER values, increased macromolecular permeability, and cytoskeletal disorganization in brain endothelial cells. Notably, these effects appear independent of changes in cell viability. Contrary to our hypothesis, plasma-derived sEVs from women with eclampsia induced less BBB impairment than those from preeclampsia or normotensive pregnancies, suggesting that soluble plasma factors—rather than sEVs—are the primary drivers of BBB dysfunction in eclampsia. This attenuated effect of sEVs may be due to reduced sEVs uptake by brain endothelial cells and decreased levels of eNOS and TNF-α within sEVs cargo. Additionally, we propose that MgSO₄ treatment may precondition eclampsia-derived sEVs, limiting their uptake and modulating their interaction with the BBB. These novel findings enhance our understanding of cerebrovascular dysfunction in eclampsia and reinforce the therapeutic relevance of MgSO₄.

To our knowledge, no prior studies have specifically investigated whether the plasma or serum of women with eclampsia directly impairs BBB integrity. However, our findings align with previous research showing that plasma from women with preeclampsia can disrupt the BBB *in vitro*^14,18,19,21,25,26^. These results reinforce the hypothesis that circulating factors contribute to a cascade of alterations, leading from BBB dysfunction to neuronal hyperexcitability, which may underlie eclamptic seizures. Notably, the absence of significant BBB disruption in cells exposed to plasma from women with complicated preeclampsia (without neurological features) suggests that eclampsia involves unique pathophysiological mechanisms beyond those observed in preeclampsia.

Our study revealed that sEVs isolated from the plasma of women with eclampsia had a weaker effect on BBB integrity than sEVs from women with preeclampsia or normotensive pregnancies. Prior studies have shown that circulating sEVs contribute to endothelial dysfunction and BBB impairment in preeclampsia^5,6,10,14^, raising the question of why this effect was less pronounced in eclampsia. These discrepancies may be attributed to differences in study populations (e.g., geographic variations: Sweden vs. South Africa), diagnostic criteria (e.g., early-onset vs. late-onset preeclampsia), or technical factors such as anticoagulants used for plasma collection (heparin vs. EDTA) or previous treatment (use or not use of MgSO_4_, see below). Additionally, differences in gestational age^27^ at sEVs isolation and storage conditions (e.g., freeze-thaw cycles) could have influenced sEV composition^28^, potentially altering their biological activity.

Moreover, O’Brien et al. demonstrated that extracellular vesicles (EVs) of unspecified size, isolated from placental explant media, did not significantly impair endothelial pro-angiogenic function. Instead, their findings suggest that soluble plasma factors, including angiogenesis-modulating proteins, may be the key contributors to vascular dysfunction in preeclampsia^29^. Then, their results reinforce the hypothesis that non-vesicular circulating components may also play a central role in BBB impairment in both preeclampsia and eclampsia.

Interestingly, our results indicate that sEVs uptake was lower in the group of women with eclampsia, significantly differing from the preeclampsia and normotensive groups. This reduced uptake may be attributed to intrinsic differences in the biophysical properties or molecular composition of sEVs from women with eclampsia. To investigate this further, we explored the role of MgSO₄, given its known endothelial stabilizing effects. Our prior work demonstrated that MgSO₄ treatment prevents BBB disruption caused by sEVs from preeclampsia-derived plasma^14^. Consistent with this, MgSO₄ significantly reduced sEVs uptake across all experimental groups. However, it did not further decrease the already diminished uptake observed in eclampsia, suggesting that early MgSO₄ exposure may influence sEVs interactions with the cerebrovascular endothelium. The exact mechanism by which MgSO₄ inhibits sEVs uptake remains unclear, but given that sEVs internalization typically occurs via endocytosis^30^, MgSO₄ may interfere with this process. Notably, women with eclampsia and most women with preeclampsia (with and without complications) received MgSO_4_ treatment. Therefore, pharmacological studies are needed to understand better how MgSO4 regulates sEVs uptake, including doses received and treatment time before delivery.

Furthermore, sEVs from women with eclampsia appear to have a distinct cargo profile compared to those from preeclampsia or normotensive pregnancies. Specifically, our analysis revealed that eclampsia-derived sEVs contain lower levels of eNOS and TNF-α, both of which are critical regulators of endothelial function and BBB integrity^31–33^. These findings suggest that alterations in sEVs cargo composition may contribute to their reduced impact on BBB dysfunction. Finally, the observation that plasma—but not sEVs—induced significant BBB damage in eclampsia suggests that additional circulating factors might be required to potentiate the harmful effects of sEVs on the cerebrovascular endothelium.

While our study provides important insights, certain limitations must be considered. First, although widely validated, our *in vitro* BBB model may not fully replicate the complex cerebrovascular dynamics observed *in vivo*. Second, prior MgSO₄ treatment in women with eclampsia and preeclampsia presents a potential confounding factor, requiring further investigation. Future research should include stratified analyses to distinguish the direct effects of MgSO₄ from intrinsic differences in sEVs composition and function

In conclusion, our study highlights the critical role of circulating plasma factors in disrupting BBB integrity during eclampsia. While plasma-derived sEVs have been implicated in endothelial dysfunction, our findings suggest that they are not the primary mediators of BBB impairment in eclampsia, likely due to lower cellular uptake and altered cargo composition. Additionally, our data support the hypothesis that MgSO₄ treatment may precondition sEVs, reducing their uptake and mitigating their impact on BBB integrity. These findings provide valuable insights into the mechanisms underlying cerebrovascular complications in eclampsia and reinforce the therapeutic importance of MgSO₄

### Clinical Perspective

Eclampsia represents the most severe cerebrovascular complication of preeclampsia, affecting approximately 1 in 100 individuals with the disorder. Its prevalence is disproportionately higher in low-resource settings, with rates ranging from 50 to 151 cases per 10,000 deliveries in regions such as Latin America and sub-Saharan Africa^1^. Despite its high morbidity and mortality, no reliable biomarkers currently exist to identify women at risk of developing eclampsia. In addition to its acute life-threatening complications, eclampsia is associated with long-term neurological consequences, including subclinical brain infarcts^34,35^, increased risk of cognitive decline^36–38^, epilepsy^39^, stroke^40^, and even dementia^41^. Early identification of at-risk women could not only improve pregnancy outcomes but also serve as a preventive strategy against future cerebrovascular disease.

A major challenge in maternal health research is the limited understanding of eclampsia’s cerebrovascular pathophysiology, which significantly impedes biomarker discovery and risk stratification. The placenta is believed to play a central role in the cerebrovascular disturbances observed in preeclampsia and eclampsia; however, its precise contribution to brain complications remains unclear. Several circulating factors—including proinflammatory cytokines, vascular modulators^18,26,32^, and placenta-derived sEVs^12,14^— have been implicated in BBB dysfunction observed in preeclampsia, yet their specific mechanistic role in eclampsia remains poorly defined.

The absence of significant BBB disruption in cells exposed to plasma from women with complicated preeclampsia (without neurological manifestations) observed in our study, suggests that eclampsia involves distinct or additional pathophysiological pathways. This insight underscores the need for targeted therapeutic strategies to address the cerebrovascular dysfunction unique to eclampsia. Moreover, further research is needed to determine whether sEVs-related pathways contribute to long-term cerebrovascular risk associated with preeclampsia and eclampsia.

Our study provides new insights by demonstrating that circulating plasma factors from women with eclampsia play a critical role in disrupting BBB integrity *in vitro*. Importantly, individual response analysis within our experimental groups suggests that plasma samples with the most pronounced BBB-disrupting effects may harbor higher concentrations of specific pathogenic factors. These findings open the possibility that *in vitro* BBB models could serve as predictive tools to identify women suffering preeclamptsia at highest risk of progressing to eclampsia. Future research should aim to characterize these plasma factors, explore their potential as biomarkers, and evaluate whether BBB dysfunction in pregnancy correlates with later-life cerebrovascular disease risk

## Abbreviations

(BMI): Body mass index
BBB: Blood-brain barrier. Claudin 5. CLDN5
FITC: Fluorescein-5-isothiocyanate
hCMEC/D3: Human brain endothelial cell line
PE: Preeclampsia
PRES: Posterior reversible encephalopathy syndrome
sEVs: small extracellular vesicles, or exosomes
TEER: Transendothelial electrical resistance
TNF-α: Tumor necrosis factor-α
VEGF: Vascular endothelial growth factor
(NP): Normotensive controls
(MgSO₄): Magnesium sulfate.

## Acknowledgment

The authors would like to thank the researchers from GRIVAS Health and RIVATREM for their valuable networking support and Dr. Fidel O Castro (University of Concepcion) for their valuable scientific and technical support in the isolation of sEVs during the generation of this manuscript. And Miss. Belen Ibáñez for her help in the initial experiments presented in this study. This study was funded by Fondecyt 1240295 (Chile). LB is funded by Wallenberg Center for Molecular and Translational Medicine. The PROVE biobank, providing the samples, is funded by research grants from the Swedish Research Council, STINT, Märta Lundqvist stiftelse, Swedish Society of Medicine, SSMF, Cornell’s foundation and Jane and Dan Olssons stiftelse.

## Author contributions

CE conceptualized the manuscript. JA, FT, EE-G performed most of the experiments. HS and FT helped with specific experiments presented in this manuscript. HS and JA prepared the draft of the manuscript. MV clinical and extracellular vesicles consultant. LB was responsible for the entire cohort of patients. PTV, LB, MV critically edited the manuscript. All co-authors approved the final version of this manuscript.

## Data availability

Data is available upon reasonable request to Prof. Carlos Escudero.

## Use of IA

We used Grammarly, a typing assistant based on Artificial Intelligence, to check English texts for grammar, clarity, and engagement. After using this tool/service, the author(s) reviewed and edited the content as needed and take(s) full responsibility for the content of the publication.

